# Competitive binding and geometric changes allow fast, complete translocation of intact HIV-1 capsids through the nuclear pore complex

**DOI:** 10.1101/2025.08.11.669633

**Authors:** Bhavya Mishra, Roya Zandi, Ajay Gopinathan

## Abstract

Recent experiments on HIV-1 capsid translocation through the nuclear pore complex (NPC) have demonstrated the docking of intact or nearly intact capsids at the pore, followed by translocation, with capsid disassembly occurring only within the nucleus near the site of integration. Given that the size of the capsid is comparable to the pore dimensions, these new findings raise questions regarding the energetics and dynamics of capsid passage across the significant entropic barrier created by the disordered FG nucleoporins (FG nups) in the NPC central channel. Here, we develop an analytical model for the transport of the HIV-1 capsid that considers the geometry of the capsid and pore, the free energy barrier due to the FG nups, and capsid-nup interactions. Our results show that capsid entry into the pore is favorable if the narrow end enters first, consistent with experimental observations, and that increasing the capsid-nup interaction strength enhances inward capsid flux. However, capsid-nup interaction alone is insufficient for complete capsid nuclear import and competitive binding by the nuclear factor CPSF6 to FG-binding sites on the capsid can serve as a ratcheting mechanism for full nuclear import. We show that nuclear import is arrested below a minimum CPSF6 concentration, consistent with experimental observations, and that pore dilation or capsid deformation can accelerate the translocation process. Our work explores the physics underlying a new perspective on HIV-1 viral genome nuclear import and provides quantitative explanations for experimentally observed phenomena while making testable predictions.

**Significance statement:** HIV-1 remains a devastating global health threat, with millions of new infections each year. However, the mechanism by which the viral genome enters the host nucleus is still not fully understood. Recent experiments show that the HIV-1 capsid can remain intact - or nearly so - during nuclear import, challenging long-held assumptions about the limits on size for transport through the nuclear pore. Here, we present a biophysical model that explains the directional transport of the capsid, the role of capsid and pore geometry and host factors such as CPSF6, and elucidates a ratcheting mechanism that allows entry of intact capsids within physiological timescales. This work provides a mechanistic framework for a critical step in the early stages of HIV-1 infection.

## Introduction

Viruses have evolved a variety of mechanisms for infecting a host cell and replicating within it [59, 77]. In cases such as HIV-1, the transport of viral genetic material into the nucleus is considered one of the most crucial steps in the virus life cycle. [13, 17, 29]. The HIV capsid, which envelops and protects both the genomic RNA and essential replication enzymes, namely reverse transcriptase and integrase, is built from 1000-1500 copies of the capsid (CA) proteins [75]. The capsid forms a conical structure with a height of about 110 nm and diameter of approximately 18 nm at the narrow end and 60 nm at the broader end [10, 21, 33, 36, 50, 60]. To reach the nucleus, the HIV-1 genome has to pass through nuclear membrane-embedded Nuclear Pore Complex (NPC) channels [16]. Because the diameter of the NPC central channel was thought to be smaller than the wide end of the HIV capsid, the prevailing model suggested that the capsid disassembles in the cytosol, allowing the viral genome to enter the nucleus through the NPC. However, recent experiments have measured the diameter of the NPC channel in SupT1-R5 cells to be approximately 65, *nm* [79], slightly larger than the wide end of the HIV capsid, suggesting that intact or nearly intact capsids may be able to pass through the NPC. In fact, several recent experiments [12, 34, 65, 78, 79] have demonstrated an intact capsid docking with the NPC and its subsequent uncoating only near the NPC’s nucleoplasmic end or within the nucleus. These observations raise questions about the fundamental energetics and kinetics of HIV capsid transport, particularly in light of the significant steric barrier posed by the crowded and narrow NPC central channel.

Traffic through the central channel is regulated by a group of nuclear pore proteins (nups) containing multiple phenylalanine-glycine repeats (termed FG-nups) [2]. These intrinsically disordered proteins, which behave as polymer chains with high conformational entropy, are grafted to the inner surface of the NPC channel [28, 40, 66, 69, 72]. FG-nups establish a size-selective diffusive barrier for cargo transport. While small cargos (5–8 nm) can passively diffuse through the central channel, larger cargos are sterically excluded. However, the NPC can facilitate the transport of cargos up to 40nm in size when they are associated with transport factors (TFs) [23, 56] such as importin and exportin proteins, which bind to the cargo. These TFs contain binding sites for the FG repeats on the FG-nups allowing the interactions to compensate for the entropic free energy penalty upon the large cargo’s entry into the central channel [5, 18–20, 41, 58, 61, 62, 73].

Once the cargo reaches the nucleus, the enzyme RanGTP binds to the TFs, cleaving them from the cargo and regulating the directionality of nucleocytoplasmic transport. This process causes the translocation of the cargo through the NPC to become ratcheted, making it unfavorable for the cargo to move backward in the channel, increasing the likelihood of release of the cargo in the nucleus [14, 25].

Given the size-selective diffusive barrier imposed by the NPC, the observed docking of the HIV-1 capsid at the pore raises significant questions. Does the capsid exhibit a directional preference for entry into the pore? Is translocation favored when the narrow end enters first? How does the capsid overcome the steric barrier created by the NUPs in the narrow channel of the NPC? What constitutes the transport signal during capsid nuclear import, and what is the underlying ratcheting mechanism? Is pore dilation (or even cracking) or capsid deformation required for complete nuclear import? While definitive answers to these questions remain elusive, recent studies have begun to illuminate several key factors that influence this process.

These factors include the presence of three important binding sites over the HIV capsid surface: the CypA binding loop, FG-binding sites, and the tri-hexamer binding surface [39, 49, 68]. Capsids are therefore able to interact with FG-nups and other cellular factors, such as CypA protein or the cleavage and polyadenylation specific factor 6 (CPSF6), via one of these interaction sites present over the capsid surface. The cohesive interaction energy gain from the capsid interaction with FG-nups could potentially compensate for the conformational entropy loss experienced by the FG-nups due to the large excluded volume of the capsid. Several experimental studies have also shown that CPSF6 plays a significant role in capsid nuclear import. Capsid localization near nuclear baskets is observed if the CPSF6 capsid binding site is deficient or mutated (A77V mutation) or if CPSF6 is depleted within the nucleus [6, 12, 78–80]. Despite these advances, a comprehensive model that incorporates established features and explains observed phenomena satisfactorily is still missing.

In this study, we develop an analytical model that incorporates known features of the HIV-1 translocation process and consensus features of the NPC to calculate the system’s free energy profile as the capsid translocates through the NPC and enters the nucleus. We explicitly consider the conformational entropy from FG-nups, the excluded volume interactions, and capsid-nup cohesive interactions in the system’s free energy. By treating capsid translocation as a Kramers’ problem [24], we compute the mean first passage time for the capsid to fully enter the nucleus as a function of several relevant parameters. This allows us to make quantitative predictions about the translocation process, which we compare with experimental observations. Our analytical calculations show that the entry of the capsid with the wider end first is prohibited due to a significant FG-nup entropic cost. This is consistent with the experimental observation that as the HIV-1 capsid enters the pore, its narrow tip faces the NPC’s cytoplasmic end [79]. It is also consistent with recent coarse-grained (CG) molecular dynamic (MD) simulations results [26] indicating a similar role for directionality in capsid entry into the pore.

Furthermore, we find that even if the capsid enters the pore with the narrow end first, transport through the NPC is energetically favorable only until the capsid’s leading tip reaches the nuclear end of the NPC. Beyond this point, a significant energy barrier exists that needs to be crossed to achieve complete import of the capsid into the nucleus. While capsid interaction with the nuclear basket FG-nups lowers the energy barrier, it is insufficient for complete nuclear import of the capsid. Our studies reveal the necessity of a ratcheting mechanism [76], which we argue arises from competitive binding by the nuclear factor CPSF6 to FG-binding sites on the capsid - a mechanism that is distinct from the RanGTP driven cleavage of TFs at the nuclear side during regular nuclear import. Specifically, we quantify how the capsid’s nuclear import and its mean translocation time depend on CPSF6 concentration within the nucleus. We show that the translocation process slows down with decreasing CPSF6 concentration and that nuclear import is arrested below a minimum CPSF6 concentration, consistent with experimental observations. Finally we also quantify the role of pore dilation and capsid restructuring/deformation that have been recently been suggested to be involved in HIV-1 capsid import [15, 78]. We show that, while for strong enough capsid-FG-nup interactions, these processes are not necessary, they can speed up translocation significantly and may be necessary for complete translocation when the capsid-nup interactions are weaker or CPSF6 concentration is lower. Our findings highlight several crucial factors regulating the translocation of the HIV capsid and its successful complete entry into the nucleus.

## Model

There is a large body of work seeking to understand cargo transport through the NPC, including studies that focus on the nuclear-protein-created size-selective barrier [5, 9, 18–20, 22, 30, 35, 41, 45, 53, 55, 57, 58, 61–64, 71, 73, 81, 82], cutoff cargo size for passive diffusion [5, 41, 44], effect of charge and hydrophobicity of nuclear-proteins [3, 70], role of pore dilation [31, 42], and cargo interaction with the nuclear-proteins for facilitated large cargo transport [4, 5, 23, 46, 47, 54, 56, 70]

Our model for the NPC, based on prior work [5], focuses on the central channel and treats it as a cylinder of length *l*_*p*_ and radius *R*_*p*_ (Fig. 1a). The pore’s inner surface is lined with flexible polymers representing the intrinsically disordered FG-nups. We assume the FG-nups are uniformly distributed over the pore’s inner surface, separated from one another by a grafting distance *d* = 10*nm*. Each FG-nup within the central channel, is assumed for simplicity to be identical and has a contour length *l*_*nup*_ = *na*, consisting of *n* monomers of length *a* ∼ 0.86*nm*. The grafted FG-nups then experience a competition between stretching and excluded volume interactions resulting in a brush of height *H* while maintaining an open central channel of radius *R*^*′*^ = *R*_*p*_ *− H* [5].

**Figure 1.**
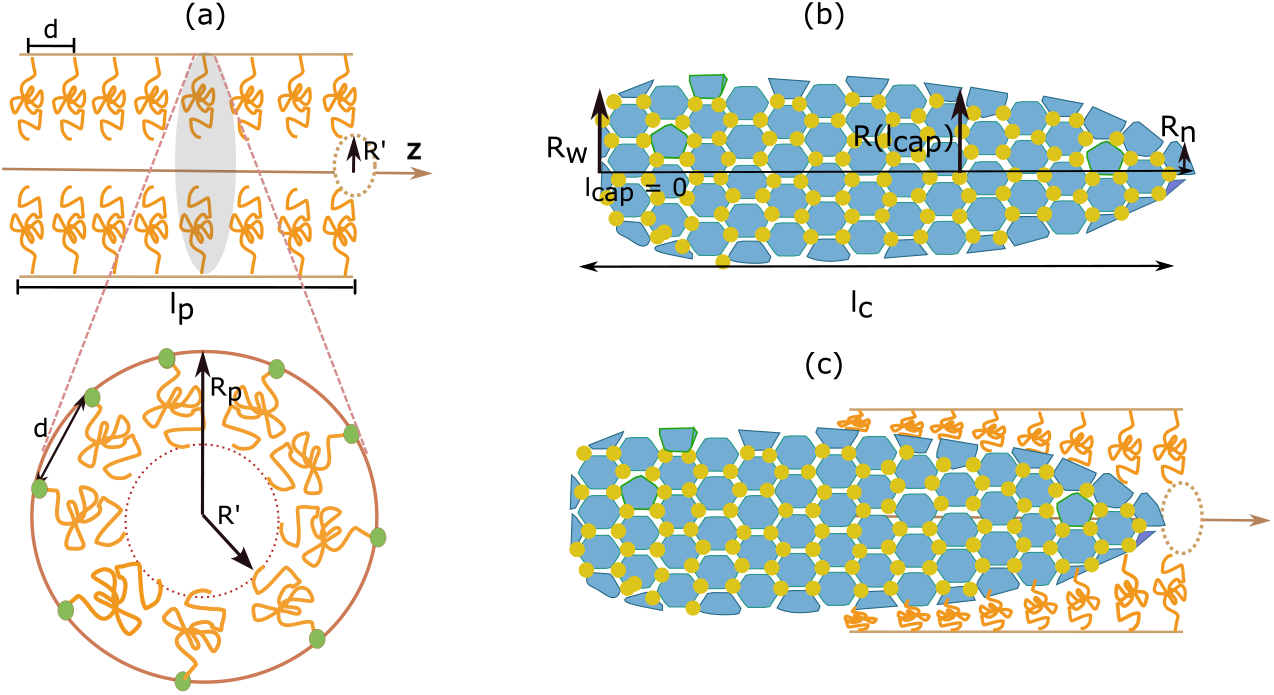
Schematic diagram of the model. (a) The NPC’s central channel modeled as a cylinder of length *l*_*p*_ and radius *R*_*p*_. FG-nups are grafted over the cylinder’s inner surface, separated from one another with a grafting distance *d*, forming a brush. (Bottom) A cross-sectional view of the cylinder: in each cross-section, *N*_*FG*_ nups are evenly distributed where green dots represent the grafting points, and *R*^*′*^ represents the central open channel radius.(b) The HIV-1 capsid is a truncated cone (narrow and wide end radii *R*_*n*_ and *R*_*w*_ respectively) with a fullerene structure comprising hexamers (blue) and pentamers (not shown) of CA proteins. Each hexamer (or pentamer) has 6 FG-binding sites over the capsid surface. (c) A capsid within the pore. The central channel FG-nups interact with the capsid via binding to the FG-binding sites with an interaction strength *ϵ*_*FG−cap*_. The capsid also displaces the FG-nups toward the pore wall, changing the brush height.

The FG-nups stretching energy *F*_*s*_, as a function of the FG-nup brush height *H* (or *R*^*′*^ = *R*_*p*_ − *H*), the number of monomers *n* per FG-nup, and the length of one monomer *a*, is given by [5],

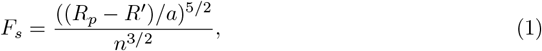

while the excluded volume interaction energy is,

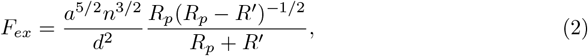

where *R*_*p*_ is the pore’s radius, and *d* is the FG-nup grafting distance. The total free energy per FG-nup can then be expressed as [5]

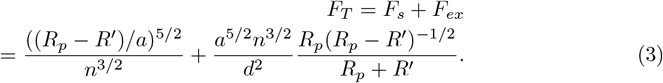

In the absence of an external perturbation within the pore, FG-nups attain their equilibrium height *H*_*eq*_, with an open central channel of radius 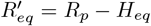 (given by by minimizing *F*_*T*_ (Eq. 3)) within the cylinder’s conduit [5]. We note that a cargo with radius greater than 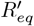 will perturb the FG-nup’s equilibrium brush height by pushing it towards the wall, lowering the FG-nup’s conformational entropy. We expect this to be a significant effect for the large HIV-1 capsid.

We now consider how the introduction of a rigid truncated cone-shaped HIV-1 capsid to our system alters the free energy. The capsid is taken to be *l*_*c*_ = 110*nm* long, with a radius of *R*_*n*_ = 9*nm* at the narrow end and *R*_*w*_ = 30*nm* at the wide end [10, 21, 33, 36, 50, 60]. The HIV-1 capsid is comprised of 250 hexamers and 12 pentamers of CA proteins separated by a distance *d*_*hex*_ ∼8*nm* [51]. The FG-nups can interact with the HIV-1 capsid via FG-binding sites that are present at the binding interface of two CA proteins within each hexamer (or pentamer) (shown as yellow dots in Fig. 1a). Since there are only twelve pentamers within the capsid, for simplicity we can take the capsid as being made of hexamers entirely as a good approximation. Now, if the capsid’s FG-binding sites are within binding range of the FG regions of the FG-nups, we expect a cohesive interaction resulting in a gain in the free-energy. We express this as (see SI),

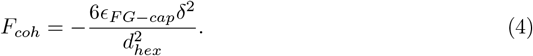

Here *ϵ*_*FG−cap*_ is the time-averaged cohesive interaction between one FG-nup and a single capsid binding site. 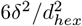 represents the average number of capsid binding sites that an FG-nup can interact with, where *δ*^2^ can be interpreted as the effective area that the FG-domains in distal region of the FG-nup cover on the capsid surface. Here we assume that *δ*^2^ scales with the dimensions of the compact hydrophobic FG-rich distal region which does not change significantly with brush conformation [73]. We also use the fact the number of FG-binding sites present over the capsid surface is always larger than the number of FG-nups available (even at the capsid’s narrow end); hence, the FG-nup number limits the maximum capsid-nup cohesive interaction.

We can now write down the free energy of the capsid-NPC system as a function of the position of the capsid’s right end, *z*, along the pore’s long axis (z-axis in Fig. 1b), with *z* = 0 denoting the capsid tip at the pore’s cytoplasmic entry. We first note that the radius of the capsid at any point *z*^*′*^ *< z* is given by a linear interpolation *R*_*cap*_(*z*^*′*^) = − (*R*_*w*_ − *R*_*n*_)(*l*_*c*_ − (*z* −*z*^*′*^)*/l*_*c*_ + *R*_*w*_ (assuming the narrow end is pointing right as in Fig. 1b). It is also to be noted that if *R*_*cap*_(*z*^*′*^) is larger than the equilibrium open channel radius 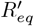, the brush is compressed to a radius *R*_*cap*_(*z*^*′*^) at *z*^*′*^ while it is left undisturbed at 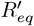 if 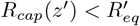. This also allows us to compute a distance *z* = *l*_*free*_ that the capsid can penetrate the pore without disturbing the brush. To compute the free energy landscapes we consider the capsid at the pore’s cytoplasmic end, *z* = 0, as the initial reference state. If the narrow end enters first, between *z* = 0 and *z* = *l*_*free*_, the free energy has only entropic contributions from the FG-nups stretching and their excluded volume interaction and is given by the reference state free energy,

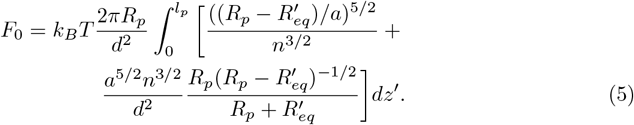

For all *z > l*_*free*_, the capsid interacts with the FG-nup brush and both entropic and enthalpic terms contribute to the free energy which we can write as,

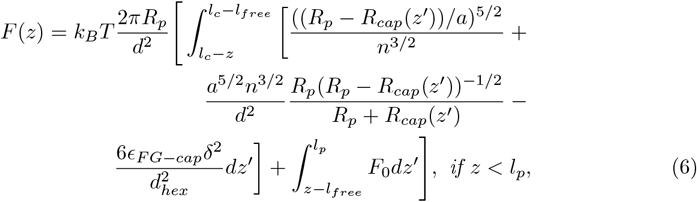

and

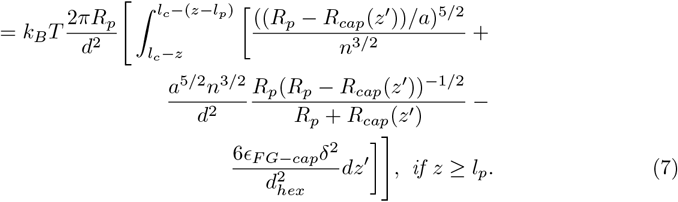

Following a similar procedure, we can derive the free energy landscape for the case where the wide end enters first.

## Results

### Capsid entry into the nuclear pore complex with the narrow end first is energetically favored

We first address the question of the existence of a directional preference in capsid translocation through the pore by studying the free energy profiles for two capsid entry modes: the narrow end entering first (narrow entry mode), and the wide end entering first (wide entry mode).

First, we define Δ*F* (*z*) as the difference in free energy between the state with capsid position at *z* (from eqs. 6 and 7) and the initial state free energy *F*_0_, i.e., Δ*F* (*z*) = *F* (*z*) − *F*_0_. We now examine the form of Δ*F* (*z*) in three different zones defined by the capsid position, *z* (see 2a). For the narrow entry mode, the capsid begins to interact with the FG-nup brush only beyond *z* = *l*_*free*_, at which point the cohesive interactions produce a decrease in Δ*F* with increasing *z*. The free energy Δ*F* reaches a minimum between zones I and II, depending on the capsid-nup interaction strength (see 2b). Beyond this point, as *z* increases, the FG-nup entropic penalty starts dominating over the capsid-nup interaction cohesive energy gain resulting in an increase in Δ*F* and a large free energy barrier. Δ*F* only begins to decrease again when the entropic gain due to the capsid leaving the pore (in zone III) overcomes the cohesive energy loss. The wide entry mode free energy profile is a mirror image of the narrow entry mode profile. It is clear from Fig. 2b that, if the wider end enters the pore first, the energy barrier that exists at the pore’s cytoplasmic end will hinder capsid import. In contrast, for the narrow entry mode, the corresponding free energy barrier only appears near the nucleoplasmic end of the pore, and capsid entry and transport through the pore are energetically favorable up to a finite distance. Increasing capsid-nup interaction strength, *ϵ*_*FG−cap*_ for the wide entry mode lowers the free energy barrier at the pore entrance (exit) but also produces a deep minimum within the pore at a distance that decreases with increasing interaction strength. Thus, from the free energy profiles, we infer that capsids are statistically much more likely to enter the pore with their narrow ends first.

**Figure 2.**
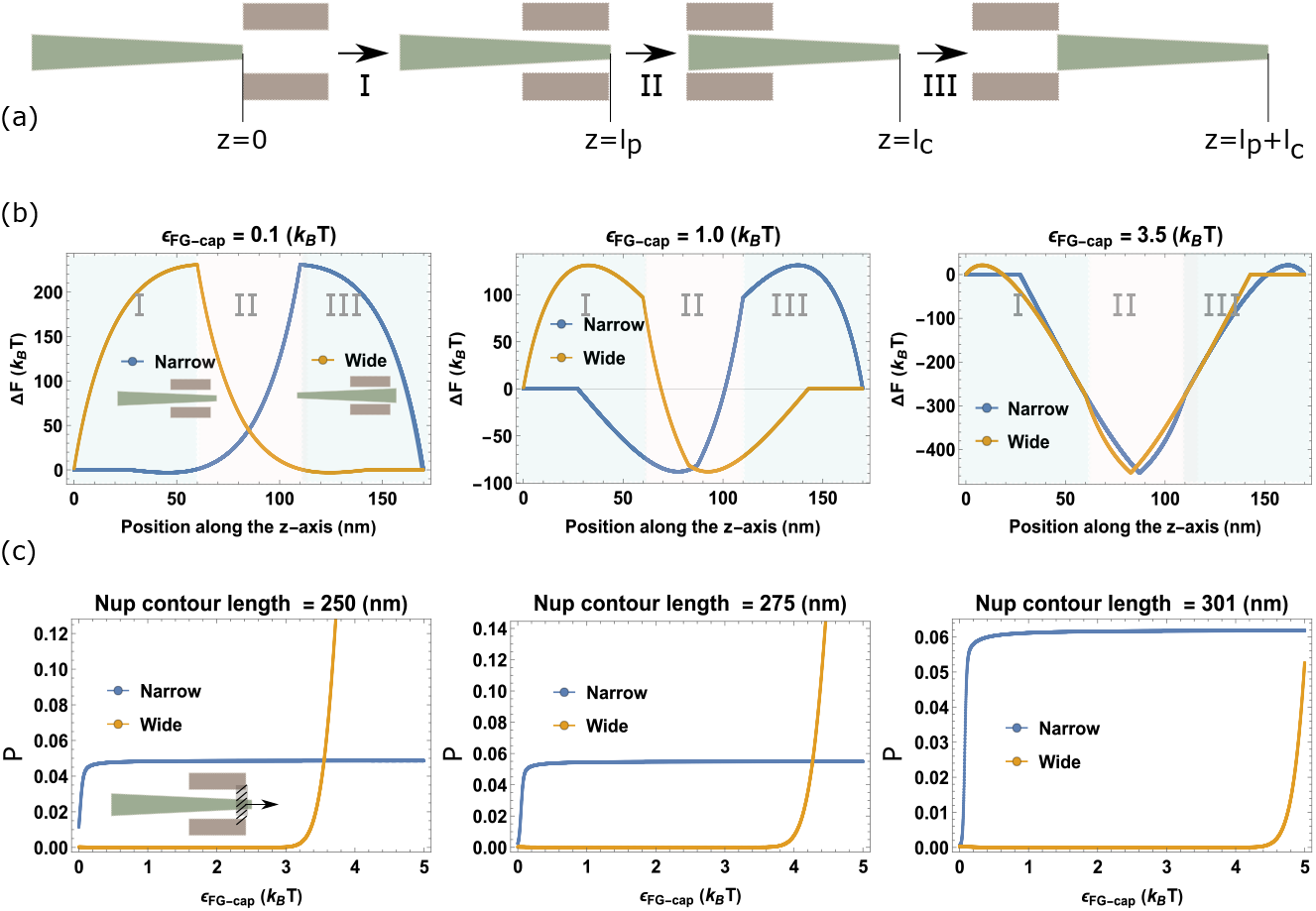
(a) Schematic representation of Zones I, II, and III indicating the position of the capsid relative to the nuclear pore. Capsid position, *z*, can be in one of three zones: Zone I spans the z values where the capsid’s right end is inside the pore, and the left end is within the cytosol, 0 *< z < l*_*p*_. In zone II, the capsid’s right end is within the nucleus, and the left is in the cytosol, *l*_*p*_ *< z < l*_*c*_. In zone III, the capsid’s right end is in the nucleus, and the left end is inside the pore, *l*_*c*_ *< z < l*_*c*_ + *l*_*p*_. (b) Comparison of the free energy profiles for narrow and wide entry modes for three different capsid-nup interaction strengths *ϵ*_*FG−cap*_ and a fixed nup contour length = 301*nm*. (c) Probability, *P*, of the capsid’s leading end entering the nucleus given that it starts at the cytoplasmic end of the pore plotted as a function of the capsid-nup interaction strength *ϵ*_*FG−cap*_ for both narrow and wide entry modes for three different nup contour lengths. For all plots, *l*_*p*_ = 60*nm, R*_*p*_ = 32.5*nm, R*_*n*_ = 9*nm, R*_*w*_ = 30*nm, l*_*c*_ = 110*nm, d* = 10*nm*, and *d*_*hex*_ = 8*nm*.

To quantify the relative probabilities of entry in the wide and narrow entry modes, we compute the probability, *P*, of the capsid’s leading end entering the nucleus given that it starts at the cytoplasmic end of the pore. This probability can be estimated by considering capsid movement as diffusion in a one dimensional potential landscape defined by *F* (*z*), and computing the probability of reaching *z* = *l*_*p*_ starting from an initial position at *z* = 0. This probability is given by [7]

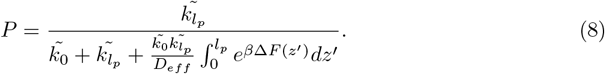

Here 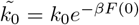 and 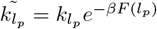, where *k*_0_ and 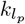 are constants characterizing the speed of capsid escape at the cytoplasmic and nuclear ends respectively. *F*_0_ is the system’s free energy when capsid is at *z* = 0, and *F* (*l*_*p*_) is the free energy at capsid position *z* = *l*_*p*_ = 60*nm. D*_*eff*_ is the effective diffusion coefficient of the capsid which is approximated to be independent of the capsid position and accounts for confinement effects due to the narrow pore (see SI). For all the numerical calculations, we take 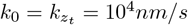 such that *k*_0_*l*_*p*_ *∼ D*_*eff*_ modeling free diffusion across the boundaries. We note that the capsid reaching the NPC’s cytoplasmic end is in itself a complex mechanism depending on a network of microtubules (MT), MT-associated molecular motors [48, 79] and a couple of cellular factors. The capsid transport to the NPC’s cytoplasmic end is beyond the scope of this work which only focuses on how the HIV-1 capsid translocates through the NPC, provided it is at position *z* = 0 at time *t* = 0.

In Fig. 2 (c) we compare the probability for the narrow and wide entry modes for varying capsid-nup interaction strengths *ϵ*_*FG−cap*_. We note that, for a fixed Nup length, the probability remains close to zero for the wide entry mode until *ϵ*_*FG−cap*_ reaches a critical value, above which wide entry mode becomes more likely. In contrast, the probability of nuclear entry is significantly higher for the narrow entry mode even for a small capsid-nup interaction strength. For reasonable values for the Nup length (*∼* 300 nm) and capsid-nup interaction strengths (*ϵ*_*FG−cap*_ ∼1 − 4*k*_*B*_*T*), we see that reaching the nucleus in the wide entry mode is extremely unlikely and docking at the cytoplasmic side with the wide end leading will almost certainly result in diffusion back to the cytoplasm. Therefore, from now on, we focus entirely on capsid entry with the narrow end first.

### Capsid stalls near the pore’s nucleoplasmic end, and the nuclear basket is insufficient for complete nuclear import

We first note that, in the narrow entry mode, the free energy is downhill till zone II, meaning the capsid can enter the pore and translocate till the narrow tip enters the nucleus. However, there is then a free energy barrier of height ∼ 100*k*_*B*_*T* that needs to be crossed for complete translocation. This implies that, without any other assisting mechanisms, complete nuclear import will be arrested, leaving the capsid trapped in the pore.

A factor that could influence this process is an additional capsid–nup interaction near the nucleoplasmic end of the pore, where the nuclear basket (NB) is located. The FG-nup NUP153, a component of the nuclear basket, is known to play a role in nuclear capsid import [11, 37]. Approximately 7–8 copies of NUP153 [38] are located at the nucleoplasmic end of the pore, where they interact with the capsid via their FG domains, which bind to FG-binding sites on the capsid with a weak binding affinity of approximately *K*_*D*_ *∼* 50 *µ*M [8]. To model the effect of NUP 153 interactions, once the capsid reaches state *z* = *l*_*p*_, we lower the free energy by 8*ϵ*_*NB−cap*_*k*_*B*_*T* where *ϵ*_*NB−cap*_ is the NUP 153-capsid interaction strength.

The modified free energy profiles in Fig. 3 exhibit a shift of 8*ϵ*_*NB−cap*_*k*_*B*_*T* in Zones II and III, reflecting the contribution of capsid–nuclear basket interactions. However, as expected from a uniform shift in the free energy landscape, the barrier to complete translocation remains of the same order of magnitude. This suggests that an additional mechanism is required to enable full nuclear import.

**Figure 3.**
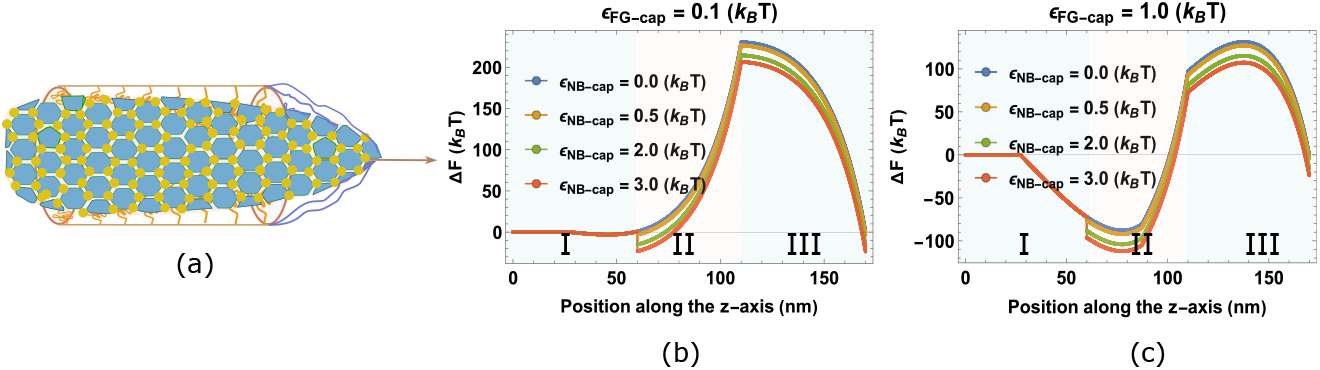
(a) Schematic representation of nuclear basket (NB), including eight copies of NUP153 at the pore’s nucleoplasmic end. The capsid segment within the nucleus binds to the NUP153 with an interaction strength *ϵ*_*NB−cap*_*k*_*B*_*T* via FG-binding sites. (b) and (c) Free-energy profiles for two different values of capsid-nup interaction strength *ϵ*_*FG−cap*_, each for five different values of capsid-NB interaction strength *ϵ*_*NB−cap*_. Parameters used: Nup length *na* = 301*nm, l*_*p*_ = 60*nm, R*_*p*_ = 70*nm, R*_*n*_ = 9*nm, R*_*w*_ = 30*nm, l*_*c*_ = 110*nm, d* = 10*nm, d*_*hex*_ = 8*nm*.

### CPSF6 capsid binding ratchets capsid transport

In addition to nuclear proteins, a key cellular factor influencing HIV infectivity is cleavage and polyadenylation specificity factor 6 (CPSF6) [1, 6, 8]. It has been shown that nuclear import can be stalled with capsids localizing near the nuclear basket if the CPSF6 capsid binding site is mutated (A77V mutation) or if CPSF6 is depleted within the nucleus [6, 12, 78–80]. This suggests a ratcheting mechanism, driven by the CPSF6-capsid binding, as a necessary condition for capsid nuclear import. In this model, the capsid surface itself serves as the nuclear transport factor containing FG-binding sites, while CPSF6 functions analogously to RanGTP in the importin-mediated translocation pathway. Rather than cleaving a host transport factor, CPSF6 competitively binds to the capsid surface, thereby preventing further interactions between the capsid and FG-nucleoporins.

To quantitatively predict the role of CPSF6 in nuclear import, it is essential to understand how CPSF6 influences the free energy profile and, consequently, the translocation times. As a starting point, we consider a uniform nuclear distribution of CPSF6 that can bind to capsid segments via FG-binding sites, provided those segments are located within the nucleus (Fig. 4a) [48, 68]. Using Langmuir adsorption kinetics, the steady-state fraction *f*_*c*_ of FG-binding sites on the capsid segment located within the nucleus that are occupied by CPSF6 can be expressed as

**Figure 4.**
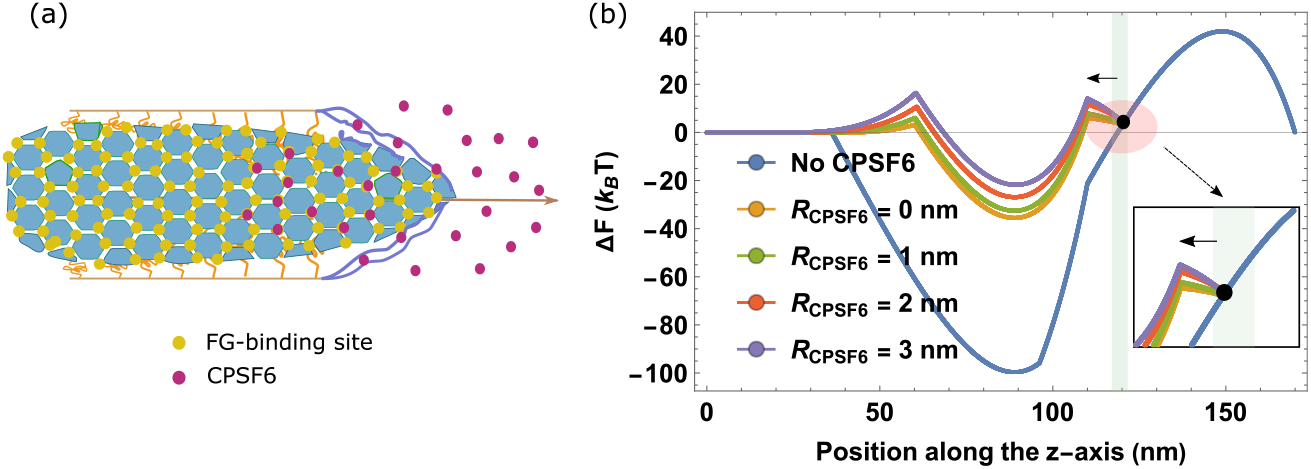
(a) Schematic representation of the nuclear basket (NB) and CPSF6 in the nucleus. CPSF6 (pink dots) binds to FG-binding sites (yellow dots) on the capsid segments that have entered the nucleus. (b) Free-energy profiles accounting for CPSF6-capsid binding for point-like CPSF6 (no entropic exclusion cost), and three different CPSF6 radii *R*_*CP SF* 6_ compared to the profile in the absence of CPSF6. If the capsid moves from position *z* (*z > l*_*p*_) (shown as a black dot) to *z*^*′*^ (*z*^*′*^ *< z*) by a distance *dz* (shown by the green band), the free energy contribution from capsid-nup interactions within the pore decreases, shifting the free energies upward. In the inset, we enlarge the free energy profiles to highlight the relative increase in total free energy upon binding of CPSF6 to the capsid.

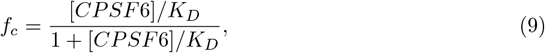

where [*CPSF* 6] denotes the concentration of CPSF6 in the nucleus, and *K*_*D*_ is the equilibrium dissociation constant (i.e., the binding affinity) for CPSF6–capsid binding, which has been experimentally measured as *K*_*D*_ ≈80 − 100*uM* [8, 52].

*We now consider how the CPSF6-capsid binding changes the free energy of the system and the capsid translocation dynamics. Once a part of the capsid is within the nucleus, CPSF6 binds to the capsid and covers a fraction, f*_*c*_, of available FG-binding sites. Now, if the capsid diffuses backwards into the pore, fewer vacant binding sites are available for FG-nups because of CPSF6 occupancy. Specifically, if the capsid starts from position *z* (with *z > l*_*p*_) and diffuses back to position *z*^*′*^ (with *z*^*′*^ *< z*), we can distinguish two regions of the capsid. The region that was never in the nucleus has an unaltered availability of FG-binding sites for capsid-nup interactions. In contrast, the part that had previously entered the nucleus has a reduced number of available FG-binding sites due to CPSF6 binding. As a result, CPSF6-capsid binding leads to a lower capsid-nup cohesive interaction in the state corresponding to *z*^*′*^. Figure 4b illustrates the change in free energy for backward diffusion from state *z* to *z*^*′*^ = *z − dz*.

If the capsid is at position *z* with *z > l*_*p*_ (shown as a black dot), CPSP6 binds to a fraction (*f*_*c*_) of FG-binding sites exposed within the nucleus. When the capsid diffuses backward by a distance *dz* to a previous position *z*^*′*^ (with *z*^*′*^ *< z*; indicated by green band), the number of capsid-nup interactions within the pore decreases. This reduction results in an increase in free energy (the yellow curve in Fig. 4b). The increased free energy at state *z*^*′*^, due to CPSF6 binding, can be expressed as a function of *z* and *f*_*c*_ by the relation, *F*_0_(*z*^*′*^) + 2*f*_*c*_*πR*_*p*_ |*F*_*coh*_| *dz/d*^2^, where *F*_0_(*z*^*′*^) is the free energy at state *z*^*′*^ in the absence of CPSF6 binding. In addition to the enthalpic contribution to the free energy, the finite size of the CPSF6 subunit that binds to the capsid can lead to an entropic penalty as it reduces the available space for the FG-nups within the pore. We can estimate this entropic contribution by replacing the capsid radius *R*_*cap*_(*z*^*′*^) in Eq.6 and Eq. 7 by an effective expanded radius *R*_*cap*_(*z*^*′*^) + *f*_*c*_*R*_*CP SF* 6_ that accounts for the size exclusion due to CPSP6 bound to a fraction (*f*_*c*_) of FG-binding sites. The total increase in free energy due to CPSF6–capsid binding effectively inhibits the capsid’s backward diffusion, thereby acting as a ratcheting mechanism that facilitates nuclear import. Note that, for the physiologically relevant parameter values chosen here, the most significant contribution to the increase in free energyrgy is enthalpic arising from the reductiincapsid-nup interactions due to CPSF6 binding (see Fig. **??** for contributions from stretching, excluded vvolume, and cohesive interactions).In what follows, we consider point-like CPSF6 to obtain a conservative estimate of the ratcheting effect while keeping in mind that the finite size of CPSF6 can additionally enhance ratcheting and lower translocation times.

To quantify the effect of the ratcheting mechanism, we calculate the translocation time when the capsid fully enters the nucleus in the presence of CPSF6. As before, we model capsid movement as diffusion in a one-dimensional potential landscape given by *F* (*z*). We consider the initial condition where the capsid’s narrow end is positioned at the cytoplasmic entry point, *z* = 0, at time *t* = 0, and we calculate the mean first passage time (MFPT) for the wide end of the capsid to reach the nuclear side, i.e. when *z* = *l*_*p*_ + *l*_*c*_. Using a reflective boundary condition at *z* = 0 (avoiding the capsid’s complete exit into the cytosol), and an absorbing boundary condition at *z* = *l*_*p*_ + *l*_*c*_ (complete translocation of the entire capsid). The MFPT can then be calculated using the standard formalism [67] as

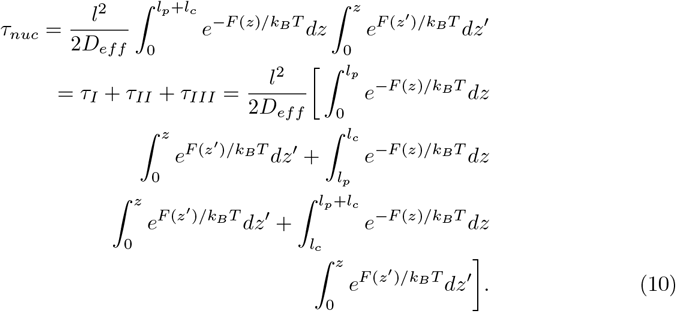

where *F* (*z*) is the free energy at state *z, F* (*z*^*′*^) is the free energy at state *z*^*′*^ and *D*_*eff*_ is the effective diffusion coefficient of the capsid. Here *τ*_*I*_, *τ*_*II*_, *τ*_*III*_ correspond to transit times in zones I, II and III respectively. We note that, since the forms of *F* (*z*) and *F* (*z*^*′*^) in the integrand of *τ*_*II*_ and *τ*_*III*_ change with *z* due to CPSF6 binding, we calculate the MFPT by numerically evaluating the integrals.

We now examine how the ratcheting action of CPSF6 influences the capsid translocation time. Figure 5 (a) shows the translocation time as a function of the normalized CPSF6 concentration, [*CPSF* 6]*/K*_*D*_, for physiologically reasonable values of the capsid-nup interaction strength, *ϵ*_*FG−cap*_ [5, 68]. *As the concentration of CPSF6 increases, the ratcheting effect becomes evident, with the capsid translocation time decreasing regardless of the strength of capsid–nup interactions. For a given value of ϵ*_*FG−cap*_, the translocation time *τ*_*nuc*_ drops sharply at low values of [*CPSF* 6]*/K*_*D*_, then gradually plateaus, approaching an asymptotic value at higher concentrations. Notably, modest changes in either *ϵ*_*FG−cap*_ or [*CPSF* 6]*/K*_*D*_ can alter the translocation time by several orders of magnitude, ranging from seconds to years.

**Figure 5.**
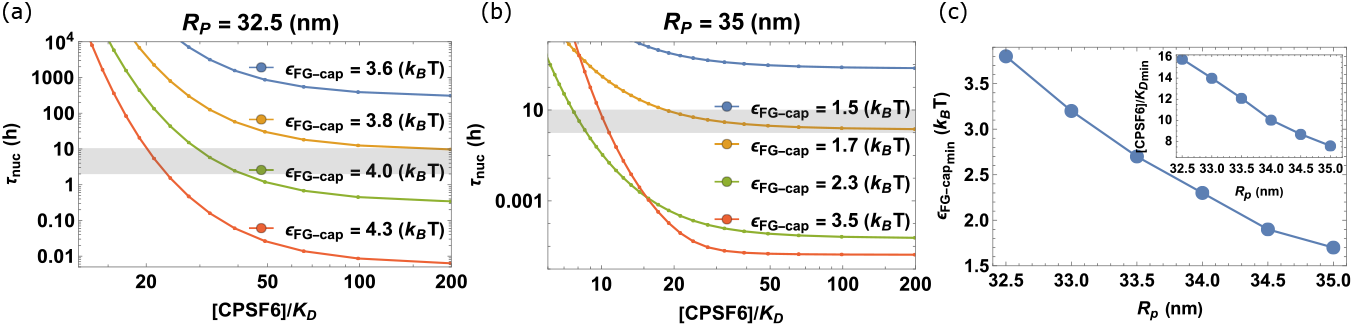
Mean first passage time (MFPT) as a function of the normalized concentration of CPSF6 for different values of capsid-nup interaction strength *ϵ*_*FG−cap*_ for pore radius *R*_*p*_ set to (a) 32.5*nm* and (b) 35.0*nm*. Horizontal gray band from 1 hour to 10 hours highlights the physiological range of time for complete nuclear import based on experimental observations [12]. (c) Minimum capsid-interaction strength *ϵ*_*FG−cap*_ required for MFPT to be lower than 10 hours at full CPSF6 coverage as a function of pore radius *R*_*p*_. Inset: Minimum normalized CPSF6 concentration needed to obtain MFPT within the physiological time range at the minimum capsid-interaction strength *ϵ*_*FG−cap*_. Parameter values used are *L* = 301*nm, l*_*p*_ = 60*nm, R*_*n*_ = 9*nm, R*_*w*_ = 30*nm, l*_*c*_ = 110*nm, d* = 10*nm, d*_*hex*_ = 8*nm, R*_*CP SF* 6_ = 0 and *ϵ*_*NB−cap*_ = 0.0*k*_*B*_*T*.

Based on experimental observations [12, 34], the physiological time scale for complete nuclear import is estimated to range from one to ten hours (indicated by the horizontal gray band in Fig.5a). For combinations of *ϵ*_*FG−cap*_ and [*CPSF* 6]*/K*_*D*_ where the mean first passage time (MFPT) exceeds this range, our model predicts that the capsid remains stalled near the nucleoplasmic end of the pore. For instance, Fig.5a shows that, given all other parameter values, capsid–nup interaction strengths below ∼3.8*k*_*B*_*T* result in stalled capsids, irrespective of CPSF6 concentration. As *ϵ*_*FG−cap*_ increases beyond this threshold, the minimum CPSF6 concentration required for physiologically relevant translocation times (i.e., under 10 hours) decreases, reaching a minimum near a critical interaction strength. However, further increases in *ϵ*_*FG−cap*_ beyond this critical point lead to a slowdown in translocation and require higher CPSF6 concentrations to achieve efficient nuclear import (see Fig. **??**).

This non-monotonic dependence arises because different regimes are dominated by different contributions to the total translocation time—namely, *τ*_*II*_ and *τ*_*III*_. When translocation accelerates with increasing *ϵ*_*FG−cap*_, the dominant barrier lies before *z* = *l*_*p*_, and the energy landscape favors forward movement once the capsid enters the nucleus. In contrast, for *ϵ*_*FG−cap*_ values above the critical point, forward movement beyond *z* = *l*_*p*_ incurs a loss in capsid–nup cohesive interactions. This loss introduces an additional free energy barrier within the nucleus, which dominates at high *ϵ*_*FG−cap*_ and results in slower translocation despite stronger initial binding, thereby requiring elevated CPSF6 concentrations to overcome the barrier and maintain physiologically relevant import times.

### NPC dilation and capsid restructuring facilitates capsid nuclear entry

Thus far, we have assumed fixed sizes for both the pore and the capsid throughout the translocation process. However, it has been shown that the nuclear envelope can undergo dilation in live cells due to mechanical strain [83]. Additionally, previous studies have identified a seam or line of defects on the HIV capsid surface [74], which could facilitate capsid restructuring. More recent evidence [79] suggests that the capsid may deform into a tubular shape during its translocation into the nucleus. We therefore now study the role of pore dilation and capsid restructuring in the HIV capsid translocation process.

We first investigate the role of pore dilation by varying the pore’s radius *R*_*p*_ while keeping other system parameters fixed. Figure 5b shows the results for a dilated pore with radius *R*_*p*_ = 35*nm*, highlighting how pore geometry influences capsid translocation. Even with a modest pore dilation of approximately 8%, we observe a dramatic drop in the MFPT by several orders of magnitude, while all other parameters remain similar (compare *ϵ*_*FG−cap*_ = 3.6*k*_*B*_*T* in Fig.5a with *ϵ*_*FG−cap*_ = 3.5*k*_*B*_*T* in Fig.5b).

Along these lines, the minimum capsid–nup interaction strength required for translocation within a physiologically relevant timeframe decreases substantially, from *ϵ*_*FG−cap*_ ∼ 3.8*k*_*B*_*T* to *ϵ*_*FG−cap*_ ∼ 1.7*k*_*B*_*T*, as the pore radius increases. We also observe a monotonic decrease in both the minimum interaction strength (Fig.5c) and the minimum normalized CPSF6 concentration (inset of Fig.5c) as the pore radius *R*_*p*_ increases from 32.5*nm* to 35.0*nm*. These results indicate that the translocation time is highly sensitive to pore geometry: even small increases in *R*_*p*_ can reduce the MFPT by several orders of magnitude and significantly lower the minimum capsid–nup interaction strength and CPSF6 concentration required for efficient nuclear import.

It is also possible that the mechanical strain induced by the entry of the capsid may itself cause the pore to dilate, as has been suggested by recent coarse grained (CG) molecular dynamics simulations [26]. Motivated by this, we wanted to understand how such a capsid position-dependent change in pore diameter might affect translocation times. To model this scenario, we fit the observed pore radius from the CG simulations to a smooth analytic function of the capsid translocation coordinate, *z*, given by *R*_*p*_(*z*) = 32.5 + 0.02*/*(0.008 + *e*^*−*0.007*z*^). We then numerically compute the MFPT as before with this *z* dependent *R*_*p*_. Note that we do not consider the energy cost for the dilation itself as we do not have a reliable estimate for it, but rather just want to get a bound for the speedup due to the change in geometry. Figure 6 compares the translocation time for this case (case 3), in which the pore undergoes dilation during capsid translocation, with two scenarios involving fixed pore radii: the original radius *R*_*p*_ = 32.5*nm* (case 1) and a uniformly dilated pore with *R*_*p*_ = 33.5*nm* (case 2). We observe a substantial reduction in the MFPT for case 3 compared to case 1, and the resulting translocation time is comparable to that of the fully dilated pore in case 2. This suggests that maximum dilation occurring toward the end of the process, when the wide end of the capsid is within the pore, has the greatest impact on reducing translocation time and is sufficient to enable physiologically reasonable nuclear import.

**Figure 6.**
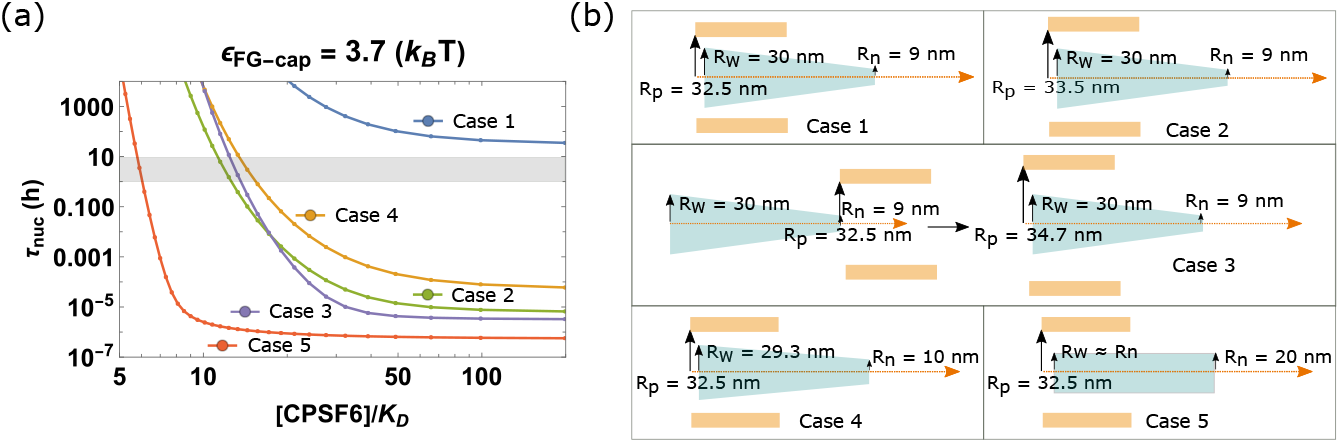
(a) Mean first passage time (MFPT) *τ*_*nuc*_ as a function of the normalized concentration of CPSF6, [*CPSF* 6]*/K*_*D*_, for five different cases of pore and capsid geometries (shown in (b)) and fixed capsid-nup interaction strength *ϵ*_*FG−cap*_ = 3.7*k*_*B*_*T* (b) Schematics of the five cases considered. Case 1: Pore radius, *R*_*p*_ = 32.5*nm*, capsid radius at the narrow and wide ends, *R*_*n*_ = 9*nm* and *R*_*w*_ = 30*nm*. Case 2: Dilated pore with radius, *R*_*p*_ = 33.5*nm* and capsid as in Case 1, *R*_*n*_ = 9*nm* and *R*_*w*_ = 30*nm*. Case 3: Pore dilation as a function of capsid position *z, R*_*p*_(*z*) = 32.5+0.02*/*(0.008+*e*^*−*0.007*z*^)*nm*. Case 4: Capsid with wider narrow end keeping volume fixed, *R*_*n*_ = 10*nm* and *R*_*w*_ = 29.29*nm*. Case 5: cylindrical capsid with the same volume, *R*_*n*_ = *R*_*w*_ = 20.4*nm*. In all cases, nup contour length *L* = 301*nm, l*_*p*_ = 60*nm, l*_*c*_ = 110*nm, d* = 10*nm, d*_*hex*_ = 8*nm, R*_*CP SF* 6_ = 0 and *ϵ*_*NB−cap*_ = 0.0*k*_*B*_*T*.

The reported flexibility of the HIV capsid [15, 43] and suggestions of restructuring during transport [79] indicates that geometric deformation of the capsid might have similar effects to pore dilation. To study the role of capsid restructuring, we change the capsid’s radius at both the narrow and wide ends while keeping the capsid’s volume and all other system parameters fixed. In 6, we increase the narrow end radius, *R*_*n*_, to 10*nm* while decreasing wide end radius, *R*_*w*_, to 29.29*nm* for case 4 and for case 5, we consider the capsid as a cylinder with an equal radius *R*_*n*_ = *R*_*w*_ = 20.4*nm* at both ends. We note that deformation or restructuring of the capsid toward a more cylindrical structure significantly speeds up translocation with the completely cylindrical capsid showing a drop in MFPT of more than *seven orders of magnitude* for saturating CPSF6 concentration. Taken together, these results suggest that geometric changes, whether in the pore, the capsid, or both, during translocation can dramatically accelerate the process. Such changes may be necessary when other parameters, such as capsid–nup interaction strength or CPSF6 concentration, fall within regimes that do not support physiologically reasonable transport times.

## Conclusion

We have developed an analytical model for the translocation dynamics of the HIV-1 capsid through the nuclear pore complex. By incorporating the geometry of both the capsid and the pore, steric constraints, specific interactions between nucleoporins and the capsid, and competitive binding by cellular factors, the model yields quantitative predictions that align with existing experimental observations and offers testable predictions for future studies.

We first quantified the extent to which the capsid entry into the pore and nucleus is energetically favorable if the capsid’s narrow end enters first. We showed that the narrow end entry mode was overwhelmingly more likely to lead to successful entry while a wide entry mode encounters a large free energy barrier and most likely results in diffusion back into the cytoplasm. Our results are consistent with experimental observations [79] of stalled capsids within the pore and those in early stages of uncoating within the nucleus all being in the narrow end first orientation. It is to be noted that we only considered interactions with FG-nups within the pore. It is however possible that capsid interactions with the cytoplasmic filament nups can also play a role in the orientation of docking at the cytoplasmic end. However, our results indicate that the orientation preference will be preserved even in the absence of cytoplasmic filaments.

Next we showed that entry via the narrow end alone is insufficient for nuclear import; an additional ratcheting mechanism, potentially provided by CPSF6-capsid binding, is essential for complete capsid nuclear import. In particular, we predict a minimum required amount of CPSF6 binding for translocation to occur. Our predictions are consistent with experimental observations showing that translocation is stalled with capsids localized near the nuclear basket when the CPSF6 capsid binding site is mutated (A77V mutation), when CypA binds competitively for the CPSF6 binding site or when CPSF6 is depleted within the nucleus [6, 12, 27, 79, 80]. We note that the ratcheting mechanism changes the interactions between the capsid and pore in a *history dependent* manner since the fraction of available binding sites for FG-nups at any point in time depends on how much of the capsid entered the pore at any time prior. Our approach is likely useful in other history dependent translocation processes that could arise in the transport of very large cargo including potentially other viruses.

We also show that, for any given set of parameters, there exists an optimal range of capsid–nup interaction strengths that enables nuclear import within physiologically relevant timescales. Direct measurements of the binding affinity of capsid FG-binding sites would be valuable in testing our prediction that physiological capsid–nup interactions fall within this optimal range, at least in the absence of geometric changes in the pore or capsid. If validated, it would be intriguing to consider whether, given the high mutational rates of viruses, the FG-binding properties of the capsid surface have undergone evolutionary optimization to maintain interaction strengths within this range.

Another remarkable result is that the translocation dynamics is extremely sensitive to geometry with small changes in geometry, either of the pore or the capsid, resulting in several orders of magnitude decreases in translocation time. This suggests that both dilation and capsid restructuring could be important *in vivo* and might even be necessary if the other parameters such as capsid-nup interactions and CPSF6 concentration are in regimes where import is not fast enough. Experimental evidence that supports these ideas include the observation of tubular forms of the capsid within the pore during transport [79], cracking of the nuclear pore [32, 78], and a deficiency in transport in the presence of mutations that affect the capsid elasticity [15, 78]. A study that couples the dilation of the pore and the elastic deformation or restructuring of the capsid due to the stresses they exert on each other would be illuminating.

Altogether, our work presents a novel and general framework for studying the translocation dynamics of intact HIV capsids through the nuclear pore complex. It identifies key regimes in the parameter space governing capsid–pore interactions and geometries that enable optimal transport. This framework can be generalized to the study of other large viral particles and may also inform the design of synthetic cargo for gene delivery and other therapeutic applications.

## Acknowledgments

This work was supported by University of California Office of the President UC Multicampus Research Programs and Initiatives, grant M21PR3267 (A.G.,R.Z.); National Science Foundation, NSF-CREST: Center for Cellular and Biomolecular Machines at UC Merced, NSF-EES-1547848 and NSF-EES-2112675 (A.G.); National Science Foundation, Center for Engineering Mechanobiology, grant CMMI-1548571 (A.G.); computing time on the Multi-Environment Computer for Exploration and Discovery (MERCED) cluster at UC Merced (NSF-ACI-1429783) and National Science Foundation, Pinnacles Computing Cluster (NSF-ACI-2019144); National Science Foundation RAPID, grant 2034794 (R.Z.); and National Science Foundation, NSF DMR-2131963 (R.Z.).

