## Supplementary Information (SI) for "Competitive binding and geometric changes allow fast, complete translocation of intact HIV-1 capsids through the nuclear pore complex"

### Free energy of capsid-nup cohesive interactions

First, we define the energy density of capsid-nup cohesive interactions, given that capsid-nup interactions have an energy $\epsilon_{FG-cap} k_B T$,

$$f_{coh} = \epsilon_{FG-cap} k_B T \rho_{FG-nup} \rho_{FG-bind} A_{pore} \tag{1}$$

where, $\rho_{FG-nup} = N_{FG}/A_{pore}$ is the number density of FG-nups, and $A_{pore}$ is the pore surface area where the FG-nups are grafted. The $\rho_{FG-bind}$ is the number density of FG-binding sites over the capsid surface and it is given by,

$$\rho_{FG-bind} = \frac{N_{FG-bind}}{A_{cap}}, \tag{2}$$

where, $N_{FG-bind} = 6.0 A_{cap}/d_{hex}^2$ is the number of FG-binding sites over the capsid surface, $d_{hex}$ is the inter-hexamer distance over the capsid surface, each hexamer contains 6 FG-binding sites and $A_{cap}$ is the capsid-surface area where the FG-binding sites are localized.

It gives us the capsid-nup cohesive interactions energy density as,

$$f_{coh} = \epsilon_{FG-cap} k_B T \frac{N_{FG}}{A_{pore}} \frac{6}{d_{hex}^2} A_{pore}. \tag{3}$$

Then, the free energy per FG-nup is equal to the area of one FG-nup times the capsid nup cohesive free energy density of all the interactions divided by the number of FG-nups,

$$F_{coh} = \frac{6 \epsilon_{FG-cap} k_B T \delta^2}{d_{hex}^2} \tag{4}$$

Where $\delta^2$ is the area of one FG-nup. For numerical solution we use $\delta = 3.6 nm$

**Table 1.** List of parameters, their numerical values and the corresponding references.

| Parameters | Numerical value | Reference |
|---|---|---|
| Nuclear pore length ($l_p$) | 60 nm | [12] |
| Nuclear pore radius ($R_p$) | 32.5 - 35 nm | [12] |
| Capsid length ($l_c$) | 110 nm | [4, 5, 7, 8, 11] |
| Capsid narrow end radius ($R_n$) | 9-10 nm | [4, 5, 7, 8, 11] |
| Capsid wide end radius ($R_w$) | 28-30 nm | [4, 5, 7, 8, 11] |
| FG-nup grafting distance ($d$) | 10 nm | [1] |
| Inter hexamer distance ($d_{hex}$) | 8 nm | [3] |
| Diffusion coefficient in free space ($D_0$) | $0.2um^2/s$ | [6] |
| Drag coefficient ($\kappa$) | 8.0-12.0 | [10] |
| Capsid-nup interaction strength ($\epsilon_{FG-cap}$) | $1 - 5k_BT$ | [1] |
| CPSF6-capsid binding affinity ($K_D$) | 80-100 uM | [2, 9] |

### Effective capsid diffusion constant

The effective diffusion constant, $D_{eff}$, of the capsid is a function of the effective viscosity of the surroundings and hydrodynamic interactions with the NPC wall. This hydrodynamic renormalization depends only on the ratio of capsid radius and pore radius. Since the capsid is an extended object with varying radii along the length, we define the effective capsid radius

$$R_c = (R_n + R_w)/2.0. \tag{5}$$

In [10], the authors calculate the drag coefficient of a spherical vesicle in a confined pore as

$$D_{eff} = D_0/\kappa(R_c/R_p), \tag{6}$$

where $\kappa$ is a drag coefficient and is a function of $R_c/R_p$. Assuming the geometry of the capsid is close enough to a sphere for this purpose, here, we derive a value for the capsid's $D_{eff}$, using the $\kappa$ values from [10] (Table 1), and $D_0 = 0.2um^2/s$ for the capsid's diffusion coefficient in a free-space. This gives $D_{eff} = 0.018um^2/s$ for $R_c/R_p = 0.6$.

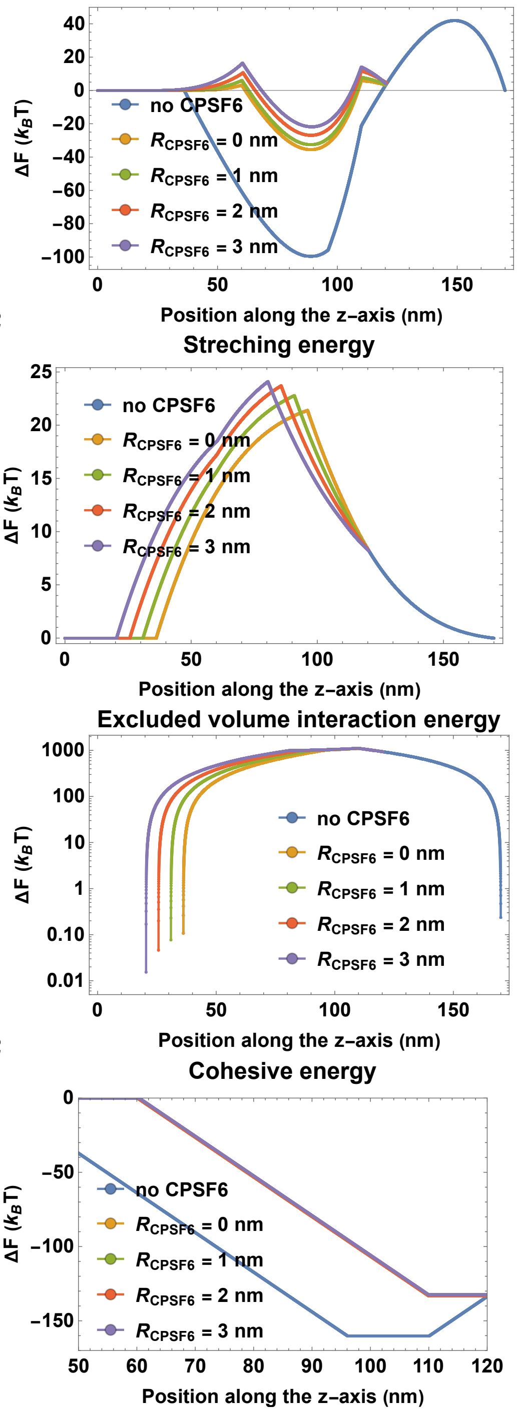


2

2

**Figure S1.** Free energy profiles for narrow mode entry for different values of the CPSF6 radius. Comparison of (top left) total free energy profiles, (top right) stretching energy, (bottom left) excluded volume interaction energy, (bottom right) capsid-nup cohesive interaction energy, in the absence of CPSF6, with point-likle CPSF6 and three different radii $R_{CPSF6}$ for a fixed nup contour length = 301nm, and capsid-nup interaction strength $\epsilon_{FG-cap} = 1.0k_BT$.

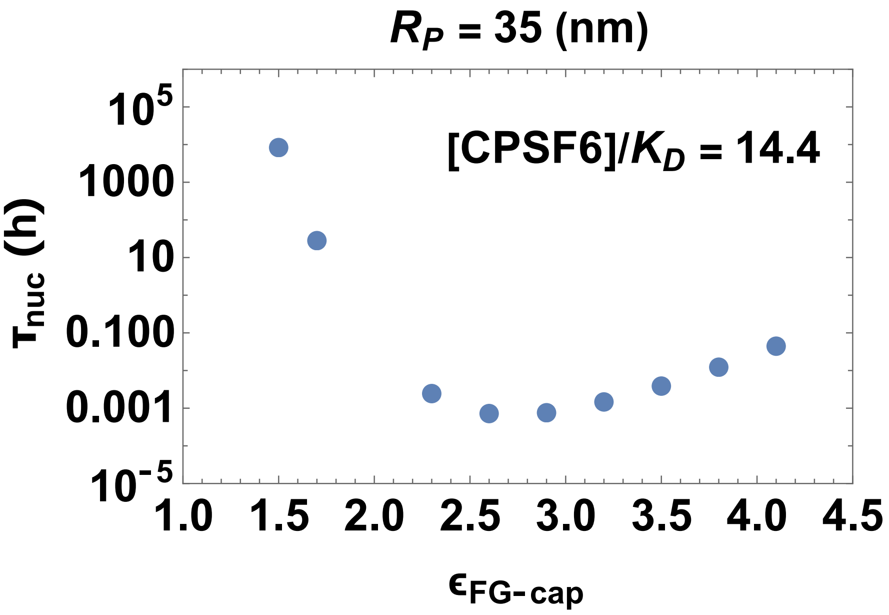


**Figure S2.** Mean first passage time (MFPT) $\tau_{nuc}$ as a function of the capsid-nup interaction strength $\epsilon_{FG-cap}$ for fixed normalized concentration of CPSF6, $[CPSF6]/K_D = 14.4$ for pore radius, $R_p = 35nm$, capsid radius at the narrow and wide ends, $R_n = 9nm$ and $R_w = 30nm$.
